# Isoform-Specific Functions of p73 Drive Survival and Chemoresistance in Diffuse Large B-Cell Lymphoma

**DOI:** 10.64898/2026.01.28.702345

**Authors:** Hesham M. Hassan, Michelle L. Varney, Dennis D. Weisenburger, Rakesh K. Singh, Bhavana J. Dave

## Abstract

Diffuse large B-cell lymphoma (DLBCL) represents 30–40% of non-Hodgkin lymphoma cases and is curable in >60% of patients; however, approximately one-third ultimately relapse. Although prior studies in normal B cells and lymphoma models implicate p73 in B-cell lymphomagenesis, the functional role of individual p73 isoforms in DLBCL remains poorly defined. TP73, a TP53 family member located on chromosome 1p36, encodes both transcriptionally active (TAp73) and dominant-negative (ΔNp73) isoforms that differentially regulate apoptosis and proliferation. In this study, we characterized p73 locus alterations, isoform-specific expression patterns, and their biological relevance in DLBCL. Chromosomal analysis revealed disruption of the 1p36 locus—predominantly via heterozygous deletion—in 35% of patient samples, which significantly correlated with elevated ΔNp73 expression. Immunohistochemical profiling demonstrated a positive association between TAp73 and cleaved caspase-3, and between ΔNp73 and Ki-67. Conversely, TAp73 expression negatively correlated with the anti-apoptotic proteins Bcl-2 and Bcl-6. Functional studies in DLBCL cell lines further confirmed that TAp73 enhances sensitivity to serum deprivation and doxorubicin, whereas ΔNp73 overexpression promotes survival and chemoresistance. Together, these findings identify p73 isoform imbalance as a key contributor to DLBCL pathogenesis and therapeutic response, highlighting ΔNp73 as a potential biomarker of aggressive disease and treatment resistance, and TAp73 as a tumor-suppressive axis warranting further investigation.

**Summary:** Diffuse large B-cell lymphoma (DLBCL) is the most common non-Hodgkin lymphoma, yet relapse remains a major challenge. The p73 gene produces two key isoforms with opposing functions: TAp73, which promotes apoptosis, and ΔNp73, which inhibits cell death and supports tumor growth. In DLBCL samples, 1p36 chromosomal disruption occurred in 35% of cases and was associated with elevated ΔNp73. TAp73 expression correlated with apoptosis markers, whereas ΔNp73 correlated with proliferation. Functional studies showed TAp73 sensitizes DLBCL cells to stress and chemotherapy, while ΔNp73 enhances resistance. These findings highlight ΔNp73 as a potential biomarker and therapeutic target in DLBCL.

## 1. Introduction

Non-Hodgkin lymphoma (NHL) is a heterogeneous group of lymphoid neoplasms with distinct morphologic, immunologic, cytogenetic, and molecular features that are associated with a specific pathogenesis for each subtype (1). Diffuse large B-cell lymphoma (DLBCL) is the most common form of NHL (about 30-40%) and is curable in more than 60% of patients (2, 3). Due to the marked heterogeneity within DLBCL, gene expression profiling has been utilized to classify DLBCL into prognostically distinct molecular subtypes (4–9). The most well-established and clinically relevant molecular subtypes are the Activated B-cell-like (ABC) DLBCL, which is associated with inferior overall survival compared with the Germinal Center B-cell-like (GCB) DLBCL (9–12). Despite long-term remission achieved with R-CHOP (rituximab, cyclophosphamide, doxorubicin, vincristine, and prednisolone), relapses occur in almost one-third of the patients (13–17). Therefore, there is a need for novel therapeutic targets that are relevant to DLBCL pathogenesis.

The *TP73* gene is a member of the *TP53* family that shares a high structural homology to p53 and is hence capable of transactivating p53 target genes (18–20). Interestingly, the *TP73* gene is located at 1p36 chromosomal locus, which is frequently rearranged in NHL (21), and disruption of the 1p36 locus was reported to be associated with a high risk of transformation of follicular lymphoma into DLBCL (9, 22, 23). The *TP73* gene locus encodes for two classes of isoforms: TAp73 (tumor suppressor isoforms retaining the transactivation domain) and ΔNp73 (oncogenic isoform, truncated and deficient in the transactivation domain), with opposing functional effects (20, 24). The balance between TAp73 and ΔNp73 isoforms and their harmony with other members of the *TP73* family regulates various cellular responses (24). Unlike p53, p73 mutations are very rare in tumors, though p73 function is frequently altered through shifting the balance between its opposing isoforms towards oncogenesis (19, 20, 25–27). The ability of p73 to substitute p53 functions, the rarity of p73 mutations, and the availability of agents that can modulate the functions of either or both isoforms make p73 a good therapeutic target in tumors.

Limited studies investigated the role of p73 in normal lymphocytes and demonstrated a potential role of p73 in the development and function of both T and B cells (28). In T-cells, it is central to T-cell activation-induced cell death (28). In B-cells, *TP73* knockout mice showed a 30% - 40% decrease in both immature and mature B cell populations in primary and secondary lymphoid organs that can be rescued by p53 co-depletion in p73^-/-^ p53^-/-^ mice (28, 29). Furthermore, high expression of ΔNp73 isoforms was found in activated and germinal center B cells, in contrast to very low levels in memory B cells and follicular mantle cells (30). Both studies suggest an important role of ΔNp73 isoforms during specific B cell developmental stages. In the lymphoma setting, using a Myc-driven B cell lymphoma transgenic mouse model, two independent studies investigated the role of p73 in the pathogenesis of lymphoma (29, 31). In the first study, a modestly significant decrease in the overall survival of p73^+/-^ mice as compared to their p73^+/+^ littermates was observed (31). In the other study, p73 did not substantially affect lymphoma onset or mortality in these mice. However, p73 loss modulated tumor phenotype in these animals in the form of increased tumor dissemination and widespread extra-nodal involvement in these mice, and this was associated with deregulation of genes involved in lymphocyte homing and dissemination (29). The constellation of these reports in normal B-cells and lymphoma animal models suggests an important role of p73 in B-cell lymphomas (26, 32). However, the biological significance of p73 isoforms in DLBCL remains unclear. Also, the feasibility of p73 modulating agents and the very low incidence of p73 mutations, combined with the importance of the p53 pathway in DLBCL, suggest that p73 could be a potential therapeutic target in DLBCL. However, little is known about the relevance of p73 to DLBCL biology and the differential role of p73 isoforms in DLBCL or other subtypes of NHL.

In this report, we performed immunohistochemical analysis for the expression of p73 isoforms and their relationship with proliferation and apoptosis in diagnostic DLBCL patient samples. We found a significant positive correlation between TAp73 and cleaved caspase-3 staining and between ΔNp73 and Ki-67. To explore the biological significance of p73 isoforms in DLBCL, we differentially modulated both TAp73 and ΔNp73 isoforms in DLBCL cell line models. TAp73-transfected cells were more susceptible to serum deprivation and doxorubicin treatment. And the opposite was seen in ΔNp73-transfected cells that were more resistant to serum deprivation and doxorubicin treatment compared to control cells.

## 2. Materials and Methods

### 2.1. Tumor specimens

Diagnostic biopsies (n = 109) of DLBCL, which were cytogenetically analyzed at the University of Nebraska Medical Center, were used for this study. The study was approved by the Institutional Review Board of the University of Nebraska Medical Center (386-15-EP).

### 2.2. Cell lines, culture conditions, and reagents

SU-DHL-16 (DHL-16) was cultured in RPMI 1640 (CellGrow™-RPMI 1640 with 2.05 mM L-Glutamine), whereas OCI-Ly3 was cultured in IMDM (CellGrow), supplemented with 10% fetal bovine serum (FBS), penicillin G (100 U/mL), and streptomycin (100 µg/mL), and maintained at 37°C in 5% CO_2_.

Over-expression was performed using HA-p73α-pcDNA3 for TAp73 and T7-p73DD-pcDNA3 for ΔNp73, and pcDNA3 for control vectors that were obtained from Addgene (Cambridge, MA) (33). Three Trilencer-27 siRNA Duplexes for *TP73* knockdown and a negative control duplex were purchased from Origene (Rockville, MD).

### 2.3. Immunohistochemistry

Immunohistochemical (IHC) analysis was performed on a tissue array prepared from diagnostic biopsies of DLBCL. The following antibodies were diluted in antibody diluent (Pharminogen, San Diego, CA) in the following ratios: anti-p73 (32-4200; Invitrogen, Grand Island, NY) which reacts with the full length human p73 (1:300); anti-ΔNp73 (IMG-313A; Imgenex, San Diego, CA), (1:1000); anti-Ki67 (H-300; Santa Cruz, Dallas, TX), (1:70); and anti-cleaved caspase-3 (ASP175 5A1E; Cell Signaling Technology, Danvers, MA), (1:200). Biotinylated secondary antibody, either anti-mouse (BA-2000; Vector Laboratories, Burlingame, CA) or anti-rabbit (BA-1000; Vector Laboratories, Burlingame, CA), diluted 1:500 in PBS and incubated for 60 min was used. Immunohistochemical evaluation by two independent observers.

### 2.4. Assessment of drug sensitivity

MTT (3-(4,5-dimethylthiazol-2-yl)-2,5-diphenyltetrazolium bromide) assay was used to determine the sensitivity and growth kinetics of the cells to serum deprivation or chemotherapeutic agents (doxorubicin or vincristine) at the indicated time points and concentrations as described earlier (33).

### 2.5. mRNA analysis

Total RNA was isolated, and real-time qRT-PCR analysis, first-strand cDNA was generated using oligo (dT)_18_ (Fermentas, Glen Burnie, MD) and Superscript II RT (Invitrogen, Grand Island, NY). 2 µL of the resulting cDNA was used in real-time PCR reactions using the primer. The primer set for TAp73 was designed to amplify all the isoforms containing the transactivation domain. For ΔNp73, the forward primer was designed to detect a sequence unique to the ΔNp73 transcript 5’ untranslated region of exon 3 (5). The primer sets for p73 proapoptotic targets BIM, PUMA, NOXA, and GRAMD4 are described earlier (33). The housekeeping β-actin gene was used to normalize gene expression. The *ct* value was normalized using β-actin for relative gene expression analysis. For the calculation of the relative expression ΔΔ *ct* was used (34).

### 2.6. Statistical analysis

The Student t-test (two-tailed) for comparisons between treated and untreated cells. One-way ANOVA test was used for comparisons between multiple groups, followed by Tukey’s and Bonferroni tests for pair-wise comparison. For correlation analysis, non-parametric bivariate correlation analyses were used (two-tailed). Chi-Square test (two-tailed) was used for comparisons between cases with or without 1p36 abnormalities in the DLBCL study. SPSS software was utilized for performing statistical analysis (SPSS Inc., Chicago, Illinois).

## 3. Results

### 3.1. TP73 gene locus (1p36) is frequently disrupted and 1p36 abnormalities are associated with higher ΔNp73 expression in DLBCL patient samples

Diagnostic samples from 109 cases of DLBCL were evaluated in this study. FISH analysis showed that 1p36 chromosomal disruption was detected in 35% of the samples. To explore whether the expression pattern of p73 isoforms correlates with 1p36 status, immunohistochemical analysis was performed for TAp73 & ΔNp73. There was no significant association between 1p36 status and TAp73 expression (**Fig. 1. A**). Interestingly, the Chi-square test showed a statistically significant association (*p* = 0.018) between ΔNp73 expression and 1p36 chromosomal disruption (**Fig. 1. A**).

**Figure 1.**
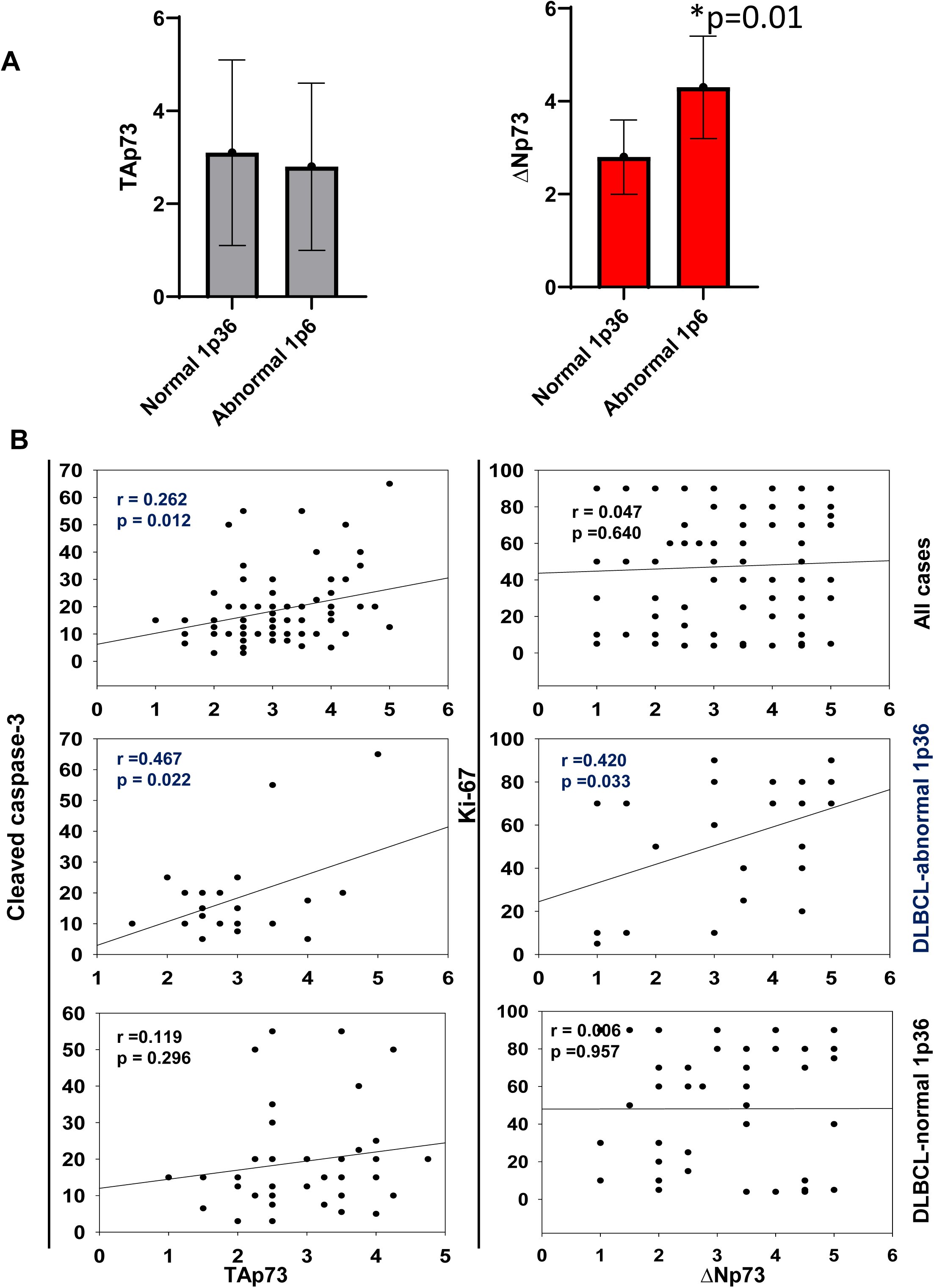
1p36 abnormalities are associated with higher ΔNp73 expression in DLBCL patient samples. **A.** Quantitative analysis of IHC scores for ΔNp73 and TAp73 expression in DLBCL biopsies (n=109) was performed. Chi-square test (two-tailed) was used to test statistical association. **B.** The differential expression of p73 isoforms correlates with apoptosis and proliferation in DLBCL patient samples (n=60). For correlation analysis, non-parametric bivariate correlation analyses (Kendall’s tau_b and Spearman’s rho) were used (two-tailed). SPSS software was used in statistical analysis.

**Figure 2.**
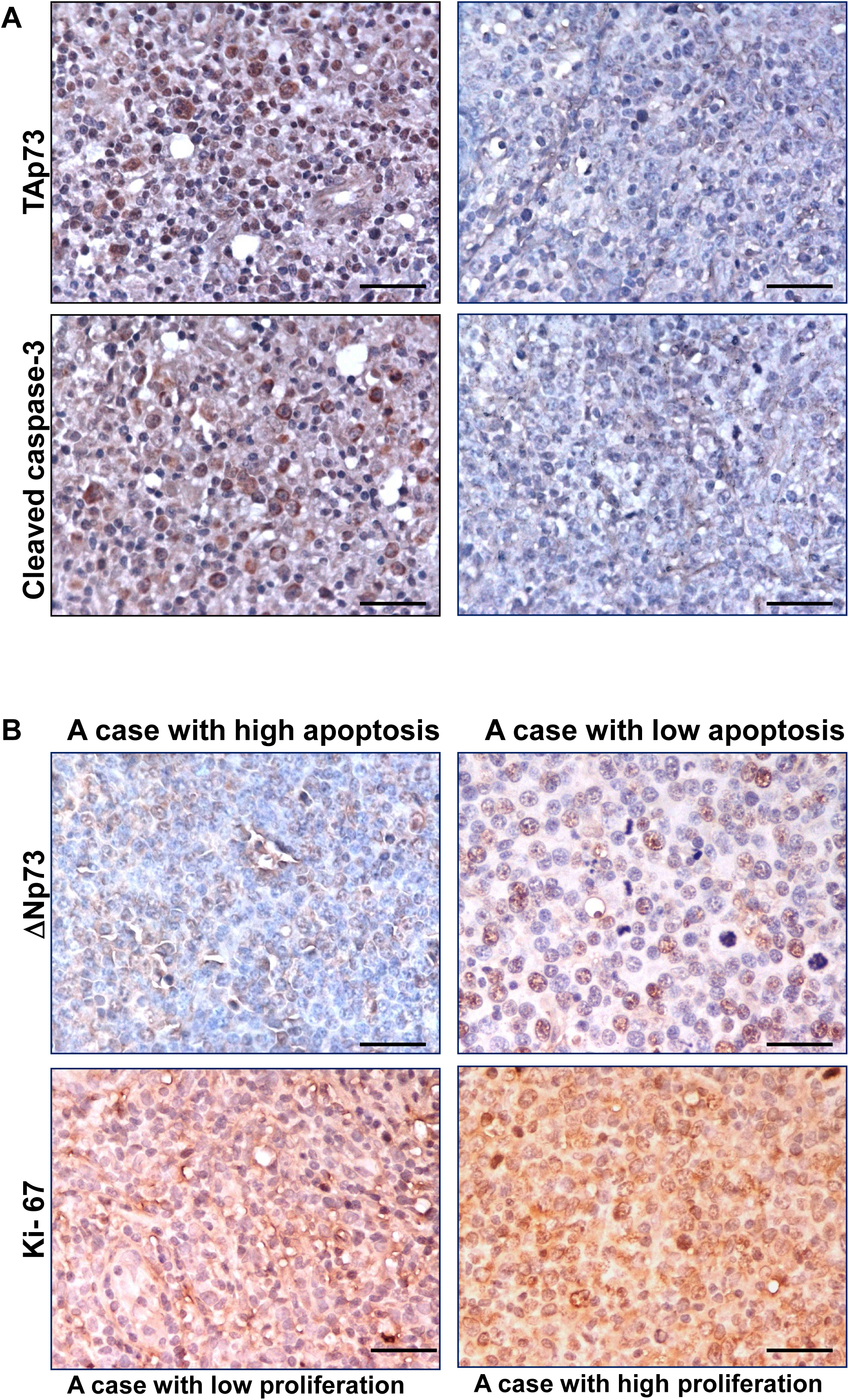
TAp73 and ΔNp73 expression in DLBCL patient samples with normal or abnormal 1p36. Representative IHC stains showing the expression of TAp73, ΔNp73, Ki-67, and cleaved caspase-3 in a DLBCL sample with abnormal or normal 1p36. The scale bar represents 50 um.

### 3.2. The differential expression of p73 isoforms correlates with apoptosis and proliferation in DLBCL patient samples

To study whether the expression of p73 isoforms correlates with apoptosis, immunohistochemical analysis was performed using cleaved caspase 3 as a marker of apoptosis. Pearson correlation analysis showed that TAp73 expression correlated significantly with cleaved caspase 3 (r = 0.262, *p* = 0.012), and that association was higher in cases that have 1p36 chromosomal disruption (r = 0.467, *p* = 0.022) (**Fig. 1B & 2A**). Furthermore, TAp73 expression was negatively associated with the expression of the anti-apoptotic protein BCL-2 (*p* = 0.04), irrespective of BCL-2 rearrangements as determined by FISH (*p* = 0.27) and BCL-6 (*p* = 0.048). To study whether the expression of p73 isoforms correlates with proliferation, the standard proliferation marker Ki-67 was used. Ki-67 staining correlated positively with ΔNp73 expression only in samples that had 1p36 chromosomal abnormality (r = 0.420, *p* = 0.033) (**Fig. 1B & 2B**).

### 3.3. TAp73 decreases the growth and increases the response of DLBCL to chemotherapy

To explore the differential effect of TAp73 isoforms on the behavior of DLBCL cells, an expression vector encoding the full-length TAp73 isoform was transfected into DHL-16 cells. TAp73-transfected DHL-16 cells showed comparable growth to control vector-transfected cells under regular conditions, as shown by MTT assay. However, TAp73-transfected cells were more susceptible than control cells to serum deprivation and the chemotherapeutic agent doxorubicin (**Fig. 3A**). Molecularly, p73 directly transcriptional pro-apoptotic targets were increased in TAp73-cells compared to control cells. Both the common p53/p73 targets (*PUMA* and *NOXA*) as well as *GRAMD4* (a p73 but not p53 pro-apoptotic target), were up-regulated in TAp73-cells compared to control cells (**Fig. 3B**). However, there was only a modest increase in the expression of the p73 target, *BIM,* in TAp73-DHL-16 cells (**Fig. 3B**).

**Figure 3.**
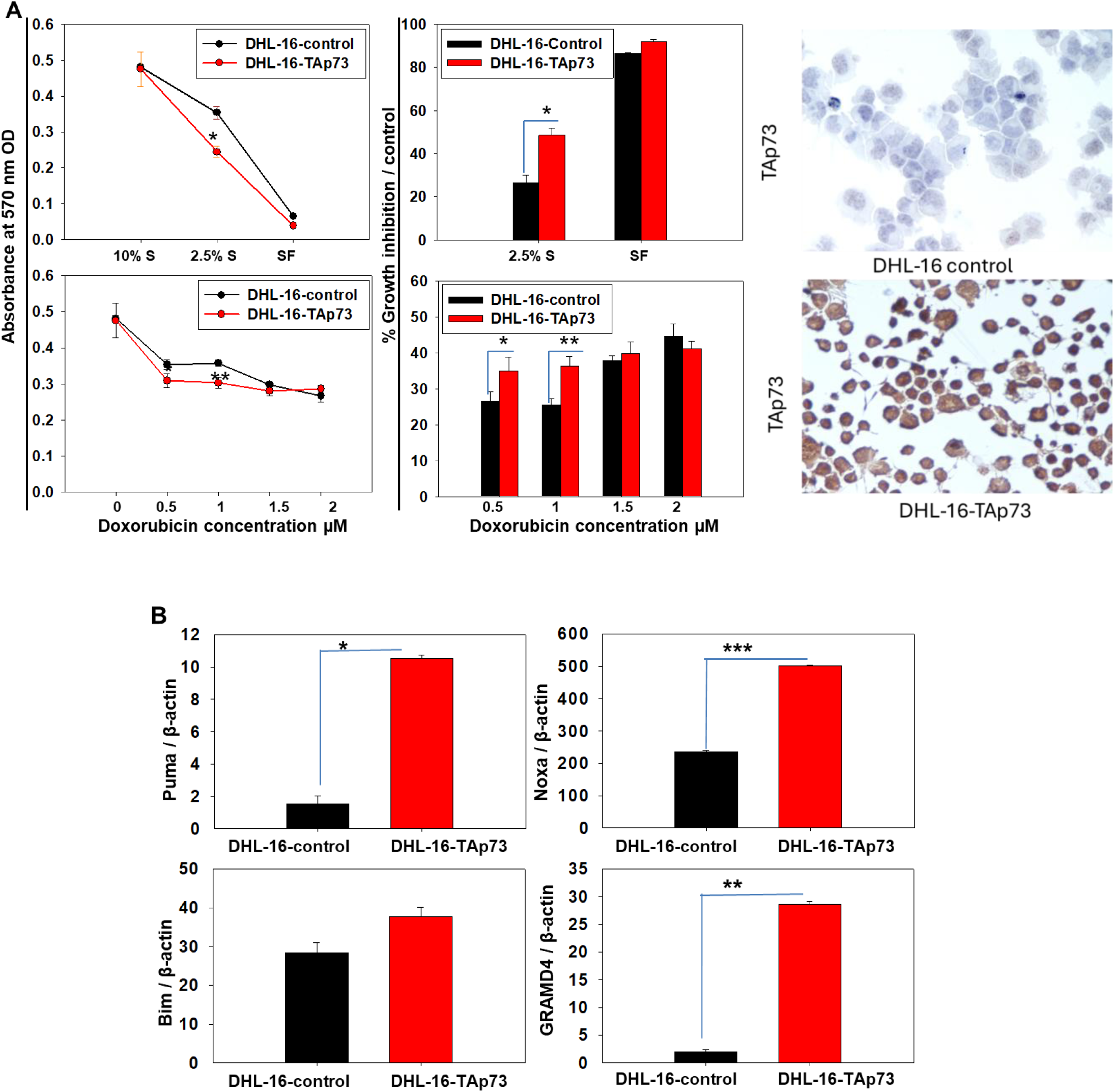
A. TAp73 decreases the growth and increases the response of DLBCL to chemotherapy. DHL-16 cells were transfected with TAp73 expression or a control vector. 48 hours after transfection, cells were grown in different serum concentrations (10%, 5%, or 0%) or treated with doxorubicin (0.5, 1, 1.5, 2 µM) for 48 hours. Then, the MTT assay was performed, and the percentage growth inhibition of the treated cells relative to their respective controls was calculated using the equation [100 - (treated/control) * 100]. * = p < .05, ** = p < .01. **B.** Increased expression of p73 transcriptional targets in TAp73-transfected cells. DHL-16 cells were transfected with TAp73 expression or a control vector. 72 hours after transfection, mRNA was isolated, followed by qRT-PCR analysis of BIM, PUMA, NOXA, and GRAMD4. * = p < .05, ** = p < .01, and *** = p < .001.

### 3.4. ΔNp73 alters the growth and cellular response of DLBCL cells to chemotherapy

To decipher the differential effect of ΔNp73 isoforms on the behavior of DLBCL cells, either an expression vector encoding a truncated protein lacking the transactivation domain similar to ΔNp73 isoforms or a control vector was transfected into DHL-16 cells. MTT assay showed that ΔNp73-transfected DHL-16 cells had an accelerated growth under regular conditions than control vector-transfected cells (**Fig. 4A**). Moreover, ΔNp73-transfected cells were more resistant than control cells to serum deprivation (**Fig. 4A**) as well as the chemotherapeutic agent doxorubicin (**Fig. 4B**). Interestingly, ΔNp73-DHL-16 cells were marginally more sensitive to the mitotic inhibitor vincristine compared to control DHL-16 cells (**Fig. 4C**). Reduction of p73 pro-apoptotic targets, *PUMA*, *BIM*, and *GRAMD4* was observed in ΔNp73-DHL-16 cells compared to control-DHL-16 cells (**Fig. 5A**).

**Figure 4.**
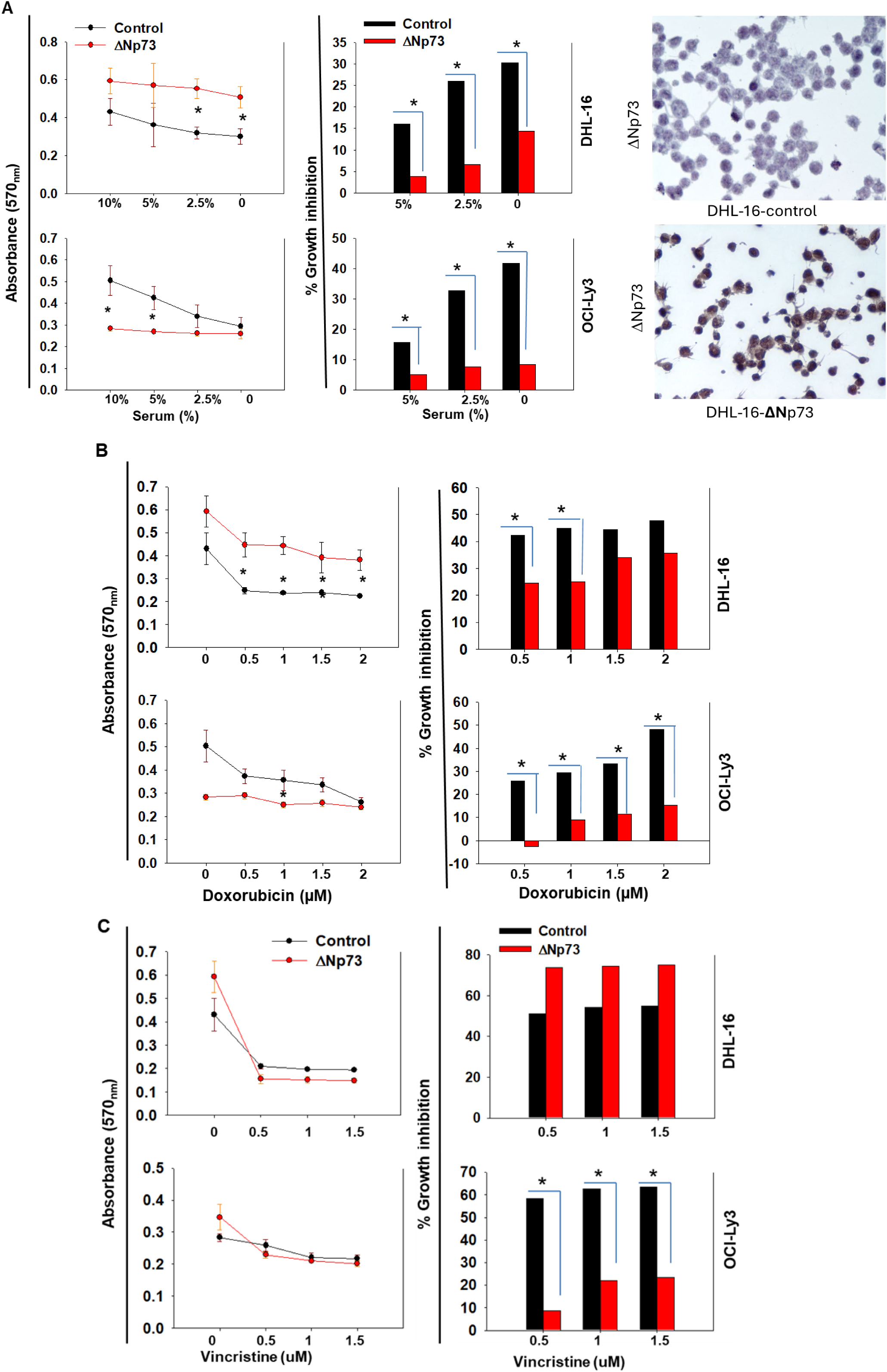
A. ΔNp73 alters the growth and the response of DLBCL cells to serum deprivation. DHL-16 or OCI-Ly3 cells were transfected with ΔNp73 expression or a control vector. 48 hours after transfection, cells were grown in different serum concentrations (10%, 5%, or 0%) for 48 hours. Then, the MTT assay was performed, and the percentage growth inhibition of the treated cells relative to their respective control was calculated with the equation [100-(treated/control)*100]. * = p < .05. **B.** ΔNp73 alters the growth and response of DLBCL cells to doxorubicin. DHL-16 or OCI-Ly3 cells were transfected with ΔNp73 expression or a control vector. 48 hours after transfection, cells were treated with doxorubicin (0.5, 1, 1.5, 2 µM) for 48 hours. Then, the MTT assay was performed, and the percentage growth inhibition of the treated cells relative to their respective control was calculated with the equation [100-(treated/control)*100]. * = p < .05. **C.** ΔNp73 alters the growth and the response of DLBCL cells to vincristine. DHL-16 or OCI-Ly3 cells were transfected with ΔNp73 expression or a control vector. 48 hours after transfection, cells were treated with vincristine (0.5, 1, 1.5 µM) for 48 hours. Then, MTT assay was performed and the percentage growth inhibition of the treated cells relative to their respective control was calculated with the equation [100-(treated/control)*100]. * = p < .05.

**Figure 5.**
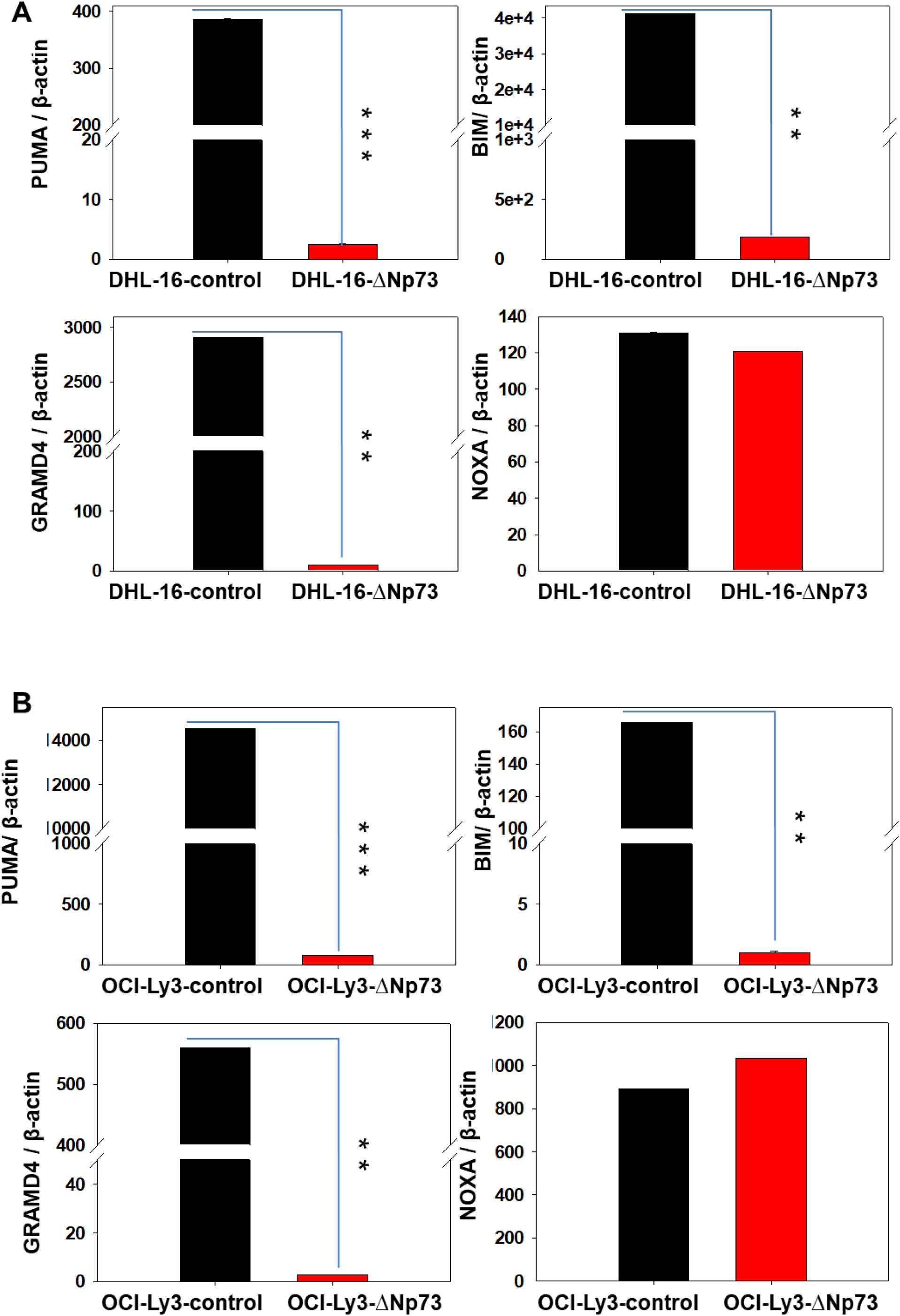
Decreased expression of p73 transcriptional targets in ΔNp73-transfected DHL-16 (A) and OCI-Ly3 (B) cells. DHL-16 and OCI-LY3 cells were transfected with ΔNp73 expression or a control vector. 72 hours after transfection, mRNA was isolated, followed by qRT-PCR analysis of BIM, PUMA, NOXA, and GRAMD4. ** = p < .01, and *** = p < .001.

Considering the well-established clinical and molecular differences between GCB and ABC molecular subtypes of DLBCL, and that DHL-16 represents the GCB subtype of DLBCL. We sought to reproduce our experiment on a DLBCL cell line of the ABC subtype, and the OCI-Ly3 cell line was used as a model of the ABC subtype of DLBCL. Surprisingly, ΔNp73-transfected OCI-Ly3 cells showed lower growth than control cells, but they were more resistant to serum deprivation (**Fig. 4A**) and both the DNA damaging agent doxorubicin (**Fig. 4B**) and the mitotic inhibitor vincristine (**Fig. 4C**). Despite the difference in behavior between the ΔNp73-transfected OCI-Ly3 and DHL-16 cells, molecularly OCI-Ly3-ΔNp73 showed reduction of p73 pro-apoptotic targets, *PUMA*, *BIM*, and *GRAMD4* similar to ΔNp73-DHL-16 cells (**Fig. 5B**).

### 3.5. p73 knockdown alters the growth and the response of DLBCL cells to chemotherapy

To further elucidate the effect of p73 isoforms on the behavior of DLBCL cells, both DHL-16 and OCI-Ly3 cells were transfected with either p73 siRNA or control siRNA. p73-siRNA-transfected DHL-16 cells showed a higher growth rate and were more resistant to doxorubicin compared to control cells (**Fig. 6A**). Conversely, there was no difference between p73-siRNA-transfected and control DHL-16 cells in the response to the mitotic inhibitor vincristine. Interestingly, p73-siRNA-transfected OCI-Ly3 cells showed a lower growth rate, but they were more resistant to doxorubicin and vincristine compared to control cells (**Fig. 6B**).

**Figure 6.**
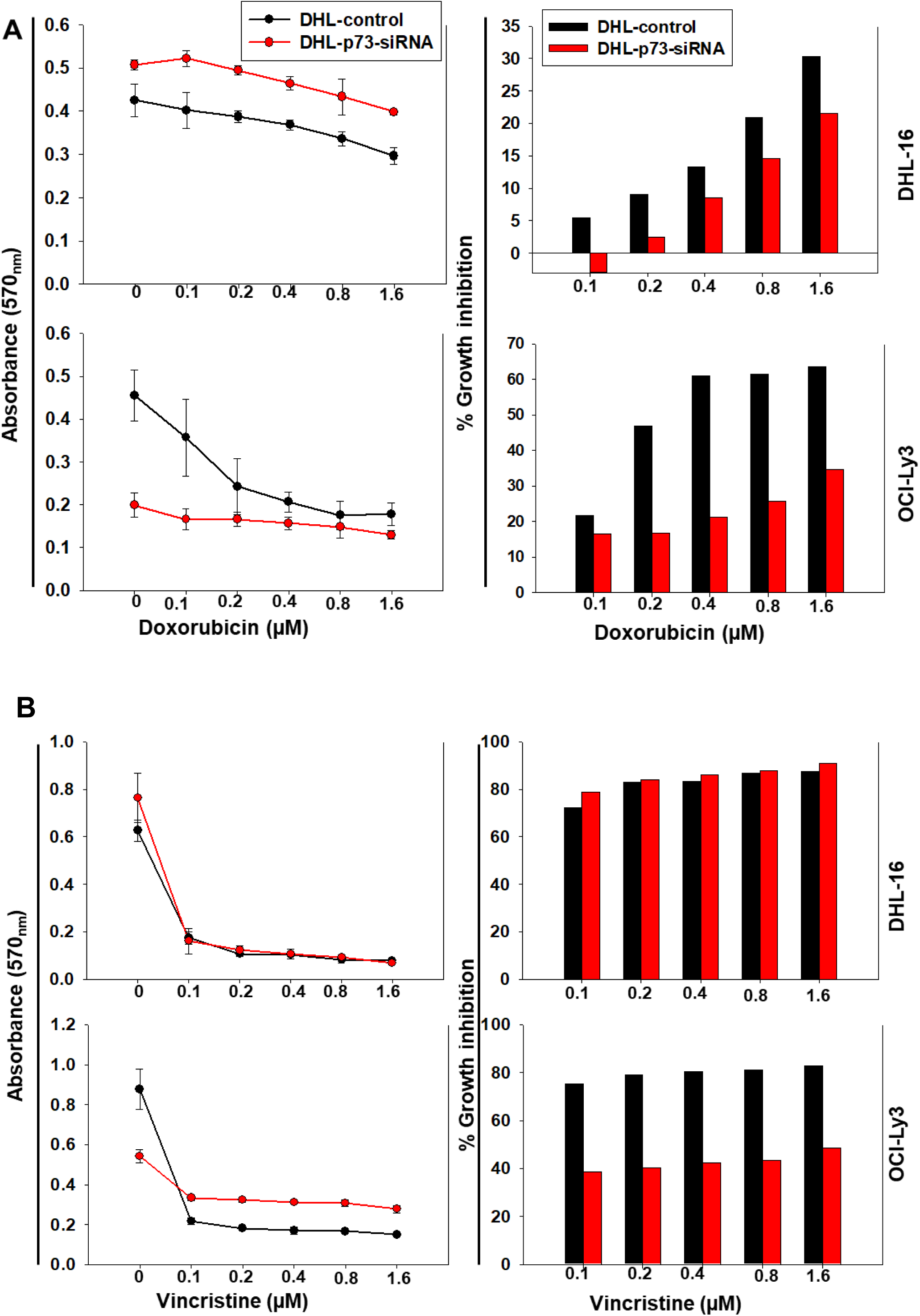
p73 siRNA alters the growth and the response of DLBCL cells to doxorubicin (A) and Vincristine (B). DHL-16 or OCI-Ly3 cells were transfected with p73 siRNA or a control vector. 48 hours after transfection, cells were treated with doxorubicin (0.1, 0.5, 1, 1.5, 2 µM) for 48 hours. Then, the MTT assay was performed, and the percentage growth inhibition of the treated cells relative to their respective controls was calculated using the equation [100 - (treated/control) * 100].

### 3.6. TP73 copy number is reduced in DLBCL, and its mRNA expression is lower in DLBCL compared to other subtypes of NHL

To further elucidate the role of p73 in DLBCL, we researched the Oncomine data sets. Interestingly, *TP73* copy number was lower in DLBCL compared to normal blood cells (**Fig. 7A**). Also, p73 mRNA expression was lower in DLBCL compared to other NHLs including the indolent follicular lymphoma as well as the aggressive Burkitt’s lymphoma suggesting peculiar relevance of p73 in DLBCL (**Fig. 7B**). Furthermore, within the DLBCL, GCB-DLBCL showed lower p73 expression compared to ABC-DLBCL subtype (**Fig. 7C**). Importantly *GRAMD4*, a novel pro-apoptotic target of p73, was lower in DLBCL compared to normal B-cells and its developmental precursors (**Fig. 7D**).

**Figure 7.**
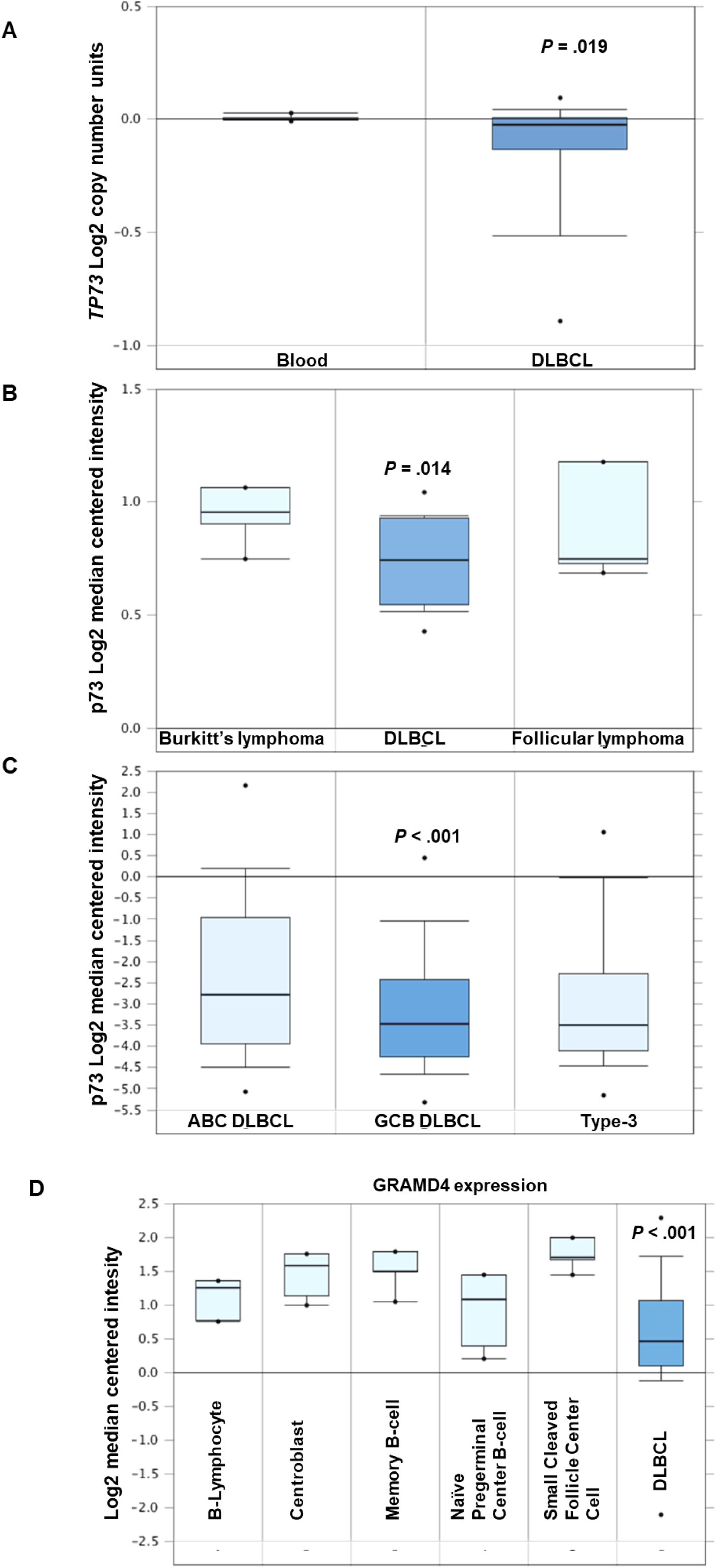
TP73 copy number is reduced in DLBCL, and its mRNA expression is lower in DLBCL compared to some other subtypes of NHL. **(A)** Copy number analysis was derived from the OCOMINE database [data was extracted from TCGA (The Cancer Genome Atlas) lymphoma]. **(B)** Analysis for p73 mRNA expression between DLBCL versus follicular and Burkitt’s lymphoma was derived from the OCOMINE database, and data were extracted from Brune et al. 2008 (35). **(C)** Analysis for p73 mRNA expression between different subtypes of DLBCL was derived from the OCOMINE database, and data were extracted from Lenz et al. 2008 (36). (D) GRAMD4 mRNA expression is reduced in DLBCL compared to normal B-cells. Analysis of GRAMD4 mRNA expression derived from the OCOMINE database, with data extracted from (37).

## 4. Discussion

In the present study, we investigated whether the modulation of TAp73 ΔNp73 isoforms expression can alter the behavior of DLBCL cell line models. In our data set, the 1p36 chromosomal locus was disrupted (mainly heterozygous deletion) in 35% of the samples and was significantly associated with ΔNp73 expression. The high incidence of 1p36 chromosomal locus deletion in DLBCL is in agreement with a previous report in which deletion of the whole short arm of chromosome 1 (1p) was detected in 28% of the total DLBCL samples included in that study (35). The relation between 1p36 chromosomal deletion and the differential p73 isoforms expression was not evaluated before in DLBCL. However, in our previous study on follicular lymphoma, there was a similar association between 1p36 chromosomal disruption and ΔNp73 isoforms expression (21, 32).

The p53 family plays a pivotal role in the regulation of many cellular biological functions, particularly DLBCL apoptosis and proliferation (36, 37). To evaluate the biological significance of p73 isoforms in DLBCL, apoptosis and proliferation were assessed using cleaved caspase 3 as a marker for apoptosis and Ki-67 as a marker for proliferation. Cleaved caspase 3 is the active form of the downstream effector caspase 3 that gets cleaved by upstream initiator caspases upon activation of the apoptotic cascade. Immunostaining for cleaved caspase 3 has been reported to be associated with a better overall survival in DLBCL (38). In our study, TAp73 expression correlated significantly with cleaved caspase-3, with a stronger correlation observed in samples harboring a 1p36 chromosomal deletion, suggesting a context-dependent role for TAp73. In contrast, Ki-67 immunostaining showed a significant positive correlation with ΔNp73 expression only in samples with 1p36 deletion, consistent with the higher ΔNp73 expression observed in these DLBCL cases. A more robust, multiparameter single-cell analysis would further strengthen and refine these observations.

To further decipher the biological significance of p73 in DLBCL, the two major classes of isoforms (TAp73 and ΔNp73) were experimentally modulated using expression vectors and siRNA. In the DLBCL cell line DHL-16, which does not express TAp73, TAp73 expression was reconstituted using a TAp73 expression vector. Interestingly, TAp73-DHL-16 cells had comparable growth under regular conditions. However, under stressful conditions such as serum deprivation or the chemotherapeutic agent doxorubicin, TAp73-DHL-16 cells were more sensitive than control cells to such conditions. The observation of enhanced effect of TAp73 over-expression upon stress can be explained by the tight regulation of TAp73 by post-translational modifications. It has been shown that chemotherapeutic agents and stress increase the stability and transactivation activity of TAp73 [reviewed in (39)]. To confirm the impact of TAp73 transfection molecularly, a panel of p73 direct transcriptional targets was measured, including *PUMA*, *NOXA*, *BIM*, and *GRAMD4*. *PUMA* and *NOXA* are well-established p53 family pro-apoptotic targets. *BIM* was reported to be a transcriptional target of p73 in regulating mitotic cell death (40) and activation-induced T-cell death (41). *GRAMD4* is a novel pro-apoptotic target of p73 that is a transcriptional target of p73 but not p53 (42). *PUMA*, *NOXA*, *BIM* (modest increase), and *GRAMD4* were up-regulated in TAp73 over-expressing cells compared to control cells, demonstrating that the phenotypic changes in TAp73-DHL-16 cells compared to control cells are mediated by TAp73 activity.

In normal B-cell development, the expression of ΔNp73 isoforms was reported to be up-regulated in activated and germinal center tonsillar B-cells. Conversely, the authors found very low levels in memory B-cells and follicular mantle cells, which suggests an important role of ΔNp73 isoforms in B-cell development (30). However, the biological significance of ΔNp73 isoforms in DLBCL was not investigated before. To gain insight about ΔNp73 isoforms in DLBCL, DHL-16 cells were transfected with an expression vector encoding a truncated protein functionally similar to ΔNp73 isoforms. ΔNp73-DHL-16 cells were more proliferating than control cells and more resistant to serum deprivation and the DNA-damaging cytotoxic drug, doxorubicin. However, there was an insignificant effect of ΔNp73 overexpression on the response to the cytostatic agent vincristine. The differential impact of ΔNp73 on response to the DNA-damaging agent doxorubicin versus the mitotic spindle inhibitory agent vincristine was predicted in a previous study (43). In that study, the authors used a kinetic modeling-based approach and in-silico cell lines to detect genetic signatures that provide chemoresistance to different classes of chemotherapeutic agents based on their mechanism of action. By pursuing their modeling strategy on in silico cell lines, they predicted that the balance between TAp73 and ΔNp73 plays a central role in a genetic signature associated with resistance to genotoxic but not cytostatic drugs (43). Furthermore, vincristine is a mitotic inhibitor that exerts its cytostatic effect through directly interacting with tubulin, disrupting the microtubules of the mitotic spindle, resulting in arrest of the cells at the metaphase of mitosis, independent of the p53 pathway (44, 45). On the contrary, doxorubicin exerts its cytotoxicity through inducing DNA damage, resulting in p53 activation and eventually cell death (46, 47). Therefore, p53 pathway activity is biologically more significant for doxorubicin cytotoxicity than the direct cytostatic effect of vincristine.

The two major molecular subtypes of DLBCL (GCB and ABC subtypes) were shown to harbor discrete genetic alterations and to have different pathogenetic mechanisms with distinct signaling pathway deregulations (35). Considering the molecular differences between the two DLBCL subtypes and that DHL-16 is a GCB subtype, we consequently sought to test the effect of ΔNp73 overexpression on a DLBCL of the ABC subtype (OCI-Ly3) (35). Interestingly, ΔNp73-transfected OCI-Ly3 cells showed slower growth but a higher resistance to serum deprivation, doxorubicin, and vincristine than control vector-transfected OCI-Ly3 cells. The difference in the phenotype observed between ΔNp73-transfected DHL-16 versus OCI-Ly3 regarding growth rate in basal conditions is expected, considering the different molecular alterations between GCB and ABC DLBCL. That explanation is supported by the observation of similar alterations in the expression of p53 pro-apoptotic target in both cell lines, despite the different phenotypes. The different responses of cell lines representing GCB and ABC subtypes to the genetic modulation of genes relevant to the molecular pathogenesis of each subtype (35). Also, ΔNp73-transfected OCI-Ly3 but not DHL-16 showed more resistance to vincristine compared to control cells. This difference in response to vincristine could be explained by the slower growth of ΔNp73-transfected OCI-Ly3 and, hence, less amenable to the cytostatic effect of the mitotic inhibitor vincristine. In contrast, the accelerated growth of ΔNp73-transfected DHL-16 renders them more liable to the cytostatic effect of vincristine.

To further elucidate the influence of p73 on the behavior of DLBCL cells, p73 siRNA was used. Interestingly, suppressing p73 activity via the siRNA approach resulted in a phenotype consistent with the phenotype observed with ΔNp73 overexpression. P73-siRNA-transfected DHL-16 cells showed higher basal proliferation and were less responsive to serum deprivation and doxorubicin but not vincristine compared to control cells. On the other hand, p73-siRNA-transfected OCI-Ly3 cells demonstrated lower proliferation, but they were less sensitive to serum deprivation, doxorubicin, and vincristine. The consistent phenotype observed with antagonizing p73 activity using either the transactivation-deficient ΔNp73 isoform or p73 siRNA suggests that the phenotype of ΔNp73-transfected cells is mainly through interfering with p73 rather than p53 activity.

## 5. Conclusion

Our studies demonstrate that the 1p36 chromosomal abnormality is frequent in DLBCL and is associated with higher expression of the dominant negative isoform of p73, ΔNp73. Biologically, ΔNp73 expression was associated with higher proliferation as evaluated by Ki-67 immunostaining in samples that harbor 1p36 chromosomal abnormalities, suggesting a more peculiar biological role of p73 in that molecular cellular context. Moreover, the expression of TAp73 (the functionally active p73 isoform) correlated with apoptosis as measured by cleaved caspase-3 immunostaining, and that correlation was stronger in samples with an abnormal 1p36 chromosomal locus, further supporting that p73 plays an imperative role in this cellular context. Experimentally, differential modulation of the p73 isoforms was associated with altered proliferation and sensitivity to serum deprivation and chemotherapeutic agents. TAp73 over-expression was associated with an increased expression of p53 family pro-apoptotic targets (*PUMA*, *NOXA*, and *GRAMD4*) and reduced basal proliferation as well as increased sensitivity to serum deprivation and the chemotherapeutic agent doxorubicin. Interfering with p73 function using ΔNp73 overexpression or p73 siRNA decreased the expression of p53 family pro-apoptotic targets (*PUMA*, *BIM*, and *GRAMD4*) and altered proliferation (increased in GCB cell line DHL-16, decreased in the ABC cell line OCI-Ly3). Furthermore, antagonizing p73 function using either ΔNp73 overexpression or p73 siRNA reduced sensitivity to serum deprivation and the cytotoxic drug doxorubicin in both cell line models and the cytostatic drug vincristine in OCI-Ly3 but not DHL-16 cells. In conclusion, the constellation of our observational and experimental observations indicates that p73 isoforms play an imperative and dynamic role in the regulation of proliferation, apoptosis, and therapy response in DLBCL. The involvement of p73 isoforms in the regulation of vital cellular activities suggests the potential of the differential modulation of p73 isoforms as a novel adjuvant therapeutic approach in DLBCL.

## Author Contributions

Writing—Original Draft: M.H.H.; Writing—Review and Editing: D.J.D. and R.K.S.; Conceptualization: M.H.H., B.J.D., and R.K.S.; Supervision: B.J.D. and R.K.S.; Experiments, acquisition of data, analysis, and interpretation: M.H.H., M.L., D.D.W., and B.J.D.; Resources and funding: M.H.H., B.J.D., D.D.W., and R.K.S. All authors read and approved the final manuscript.

## Institutional Review Board Statement

We obtained tissue microarray (TMA) slides from the Lymphoma Study Program of the University of Nebraska Medical Center (UNMC), Omaha, Nebraska (NE), in compliance with IRB# 386-15-EP).

## Informed Consent Statement

This exempt study involved no recruitment, and therefore no informed consent was required.

## Data Availability Statement

The data presented in this study are available in the article or supplementary material published.

